# Temperature-induced changes in protein interactions control RNA recruitment to G3BP1 condensates

**DOI:** 10.1101/2024.02.02.578543

**Authors:** Charlotte M. Fischer, Hannes Ausserwöger, Tomas Sneideris, Daoyuan Qian, Rob Scrutton, Seema Qamar, Peter St George-Hyslop, Tuomas P. J. Knowles

## Abstract

Biomolecular condensates have emerged as prominent regulators of dynamic subcellular organisation and essential biological processes. Temperature, in particular, exerts a significant influence on the formation and behaviour of biomolecular condensation. For example, during cellular heat stress, stress granules (SGs) are formed from RNA-binding proteins (RBPs) and RNA, forming liquid condensates to protect the RNA from damage. However, the molecular mechanisms leading to changes in protein phase behaviour are not well understood. To answer how temperature modulates protein interactions and phase behaviour, we developed a high-throughput microfluidic platform, capable of mapping the phase space and quantifying protein interactions in a temperature-dependent manner. Specifically, our approach measures high-resolution protein phase diagrams as a function of temperature, while accurately quantifying changes in the binodal, condensate stoichiometry and free energy contribution of a solute, hence, providing information about the underlying mechanistic driving forces. We employ this approach to investigate the effect of temperature changes on the phase separation of the stress granule scaffold protein Ras GTPase-activating protein-binding protein 1 (G3BP1) with PolyA-RNA. Surprisingly, we find that the G3BP1/RNA phase boundary remains unaffected by the increasing temperature but the underlying stoichiometry and energetics shift, which can only be revealed with high-resolution phase diagrams. This indicates that temperature-induced dissolution is counteracted by entropic processes driving phase separation. With increasing temperature, the G3BP1 content in condensates decreases alongside with a reduction of the free energy of protein interactions, while the RNA content increases driven by entropically favoured hydrophobic interactions. In the context of cellular heat SG formation, these findings could indicate that during heat shock, elevated temperatures directly induce RNA recruitment to stress granules as a cytoprotective mechanism by finetuning the strength of protein and RNA interactions.

## Introduction

Biomolecular phase separation has gained considerable attention in recent years due to its critical role in various biological processes^1,2^ and its implications in material science and engineering^3–6^. Phase separation involves the spontaneous demixing of a homogeneous liquid into two distinct phases, a biopolymer-rich dense phase and a biopolymer-poor dilute phase. In the context of cellular biology, phase separation is the key process governing the formation of membraneless organelles, which are essential for cellular functions such as RNA processing^7–10^, signal transduction^11–16^, and stress response^17–20^. These membraneless organelles are dynamic and undergo constant remodelling, allowing cells to quickly adapt to changing environmental conditions^21,22^. Nucleoli^23–25^ and stress granules (SGs)^26,27^, in particular, exhibit temperature sensitivity. Specifically, changes in temperature of the environment modulate the formation, dissolution, and dynamics of these organelles, affecting cellular functions and signalling pathways^23,24,26^. For instance, defective SG formation and disassembly can have implications in viral infection^28,29^, inflammatory^29–31^ and neurodegenerative disease^32–37^ such as amyotrophic lateral sclerosis (ALS) or frontotemporal dementia (FTD) as well as tumour progression^38–43^. Understanding the mechanism of temperature modulation on the molecular level is key to developing therapeutics to reverse these processes.

While the impact of temperature on cellular condensation is well documented, the precise mechanisms underlying these effects remain elusive. We aim to unravel how temperature influences protein interactions, thereby modulating phase separation phenomena. To address this question, we establish a high-resolution and -throughput approach for studying the effects of temperature on protein phase separation and collective, emergent interactions^44–47^ based on combinatorial microfluidic technology^48,49^. Using the PhaseScan^50^ platform, we precisely measured FUS and G3BP1 protein phase diagrams with multiple solute axes at different temperatures. This approach enabled us to monitor shifts in the position and change in the shape of the phase boundary, characterise protein dilute phase concentrations, and determine tie-lines and solutes’ free energy contributions, unravelling information about the changes in stoichiometry and interactions of condensate components with changing temperature.

We utilise our platform to study the temperature-dependence of G3BP1, a scaffold protein of SGs, in the presence of unstructured PolyA-RNA. Surprisingly, we find that the position of the phase boundary is barely affected by temperature as thermal dissolution is counteracted by entropically favourable processes. The stoichiometry of the components inside the G3BP1/PolyA condensates changes drastically towards higher RNA content with increasing temperature. At the same time the contribution of G3BP1 to the free energy of phase separation diminishes, while RNA interactions become the dominant driving force of phase separation at higher temperatures. We hypothesise that solute interaction changes induced by elevated temperatures cause the recruitment of untranslated RNAs into SGs as part of a cytoprotective mechanism.

## Results and Discussion

### Quantifying temperature-dependent protein phase behaviour using droplet microfluidics

To investigate the molecular mechanisms of temperature modulation of protein interactions and phase separation, we employed a combinatorial microfluidics-based platform ‘PhaseScan’.^50^ Three-dimensional phase diagrams were acquired via the platform illustrated in Fig. 1a. Briefly, droplets of different composition containing protein (FUS or G3BP1) and modulator (PEG or RNA) were generated on a chip by varying the flow rates of the aqueous solutions. The aqueous solutions of protein and modulator were barcoded using fluorescent dyes. The microfluidic chip was positioned on a heating stage and set to a constant temperature to allow for accurately varying the temperature of the droplets inside the microfluidic chip. On the way to the imaging chamber, the droplets underwent incubation at the desired temperature for 5 minutes. Subsequently, the flowing droplets were continuously imaged via wide-field fluorescent microscope in the imaging chamber.

**Fig. 1:**
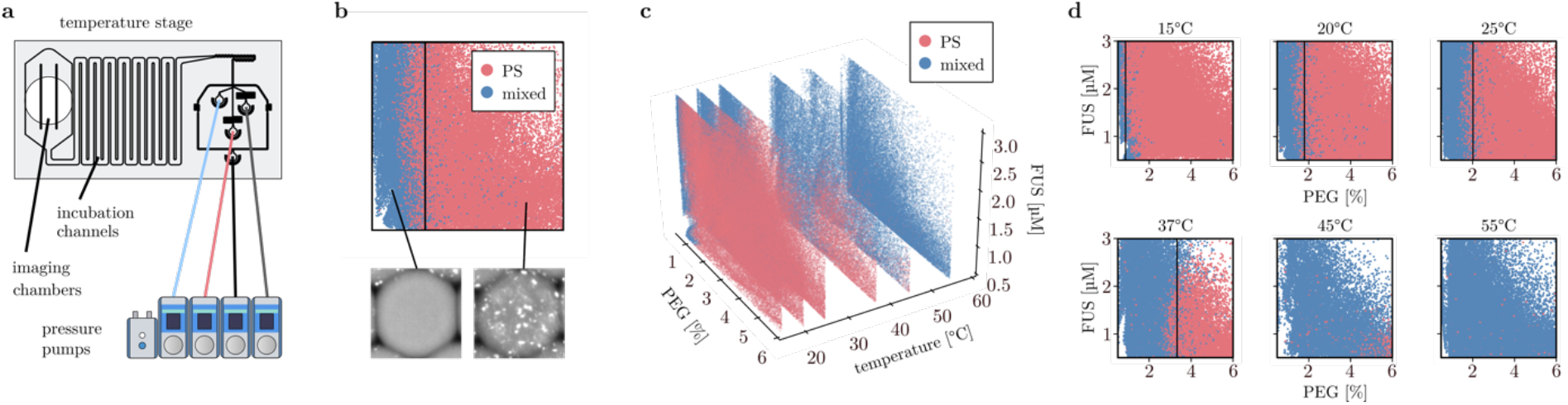
Phase separation of the FUS/PEG system at varying temperature. **a** A microfluidic chip mounted on a temperature control stage is used to perform the experiments, encompassing the standard droplet maker, incubation channels, and large imaging chambers. Condensates are detected in each droplet, using a standard epifluorescence imaging setup. **b** Total concentrations of the protein and modulator are determined from total fluorescent intensities and the phase diagram is plotted from the droplet data. **c** 3D phase diagram of FUS as a function of PEG concentration and temperature. The diagram is a combination of 2D phase diagrams acquired at 15, 20, 25, 37, 45 and 55°C. Points in the phase diagram correspond to individual droplets. N=298860 **d** Panel 1-6: Phase diagrams of FUS as a function of PEG 10K concentration. Points in the phase diagram correspond to individual droplets. The phase separated regime is represented in red, the re-entrant homogeneous phase is shown in blue. N= 61331, 66311, 74748, 30186, 18396, 47888 data points for 15 to 55°C, respectively.

A convolutional neuronal network (CNN) was utilised to process the images, detect droplets and classify them as phase separated or mixed based on the presence or absence of condensates, respectively (Fig. 1b). The fluorescence intensity of each barcode inside the droplet was measured to calculate the concentration of the labelled components. For determining the dilute phase concentration, the darkest 5-25% pixels inside the droplet were used to calculate the concentration of the protein outside the condensates. Using this data, phase diagrams, as well as tie-line and solute free energy diagrams for different temperatures could be plotted with each point in the phase diagram representing one droplet with a unique microenvironment.

First, we employed our platform to study the temperature modulation of FUS as a model system. FUS is a phase separating protein involved in nucleic acid metabolism^51^ but also linked to neurodegenerative disease^52–55^. We acquired two-dimensional phase diagrams of FUS with PEG at six different temperatures between 15 and 55°C (Fig. 1c-d). The phase boundaries of the FUS-PEG system were approximately parallel to the FUS axis in the concentration and temperature ranges investigated. Their positions were determined by fitting the data with a linear support vector machine (SVM) fit. A significant shift of the phase boundary (Fig. 1d; black dashed line) of the FUS-PEG system towards condensate dissolution was observed upon temperature increase, ranging from 0.9% PEG needed to induce phase separation in a range of 0-3µM FUS at 15°C to 6.4% PEG needed at 45°C. At temperatures higher than 55°C, no condensate formation could be observed under any of the conditions screened in this experiment. The results are in line with previous observations on the upper critical solution temperature (UCST) behaviour of FUS^56,57^, which, like most phase separating protein systems, exhibits UCST behaviour, such as Tau^58,59^, hnRNPA1/2^60,61^ or DDX4^62^, where the entropic gain from mixing outweighs the enthalpic gain from protein interactions at higher temperatures.

### Temperature of the environment modulates component stoichiometries and intermolecular interaction strength inside condensates

LLPS is driven by the free energy difference between the homogeneous and the phase separated state. This energy difference underlies the differences in concentrations between the dilute and the dense phase which can be described by a tie-line vector^45,47^ (Fig. 2a). The tie-line vector connects the dilute phase to the corresponding dense phase concentration. Therefore, measuring the tie-lines for a system at different temperatures yields information about the stoichiometric changes in the condensates. The free energy difference of the phase separating system can be interpreted as a sum of contributions by each individual solute species. For each solute, the dominance factor D quantifies the relative weight of the contribution of the interactions of a solute to the whole system^45,46^ (Fig. 2b).

**Fig. 2:**
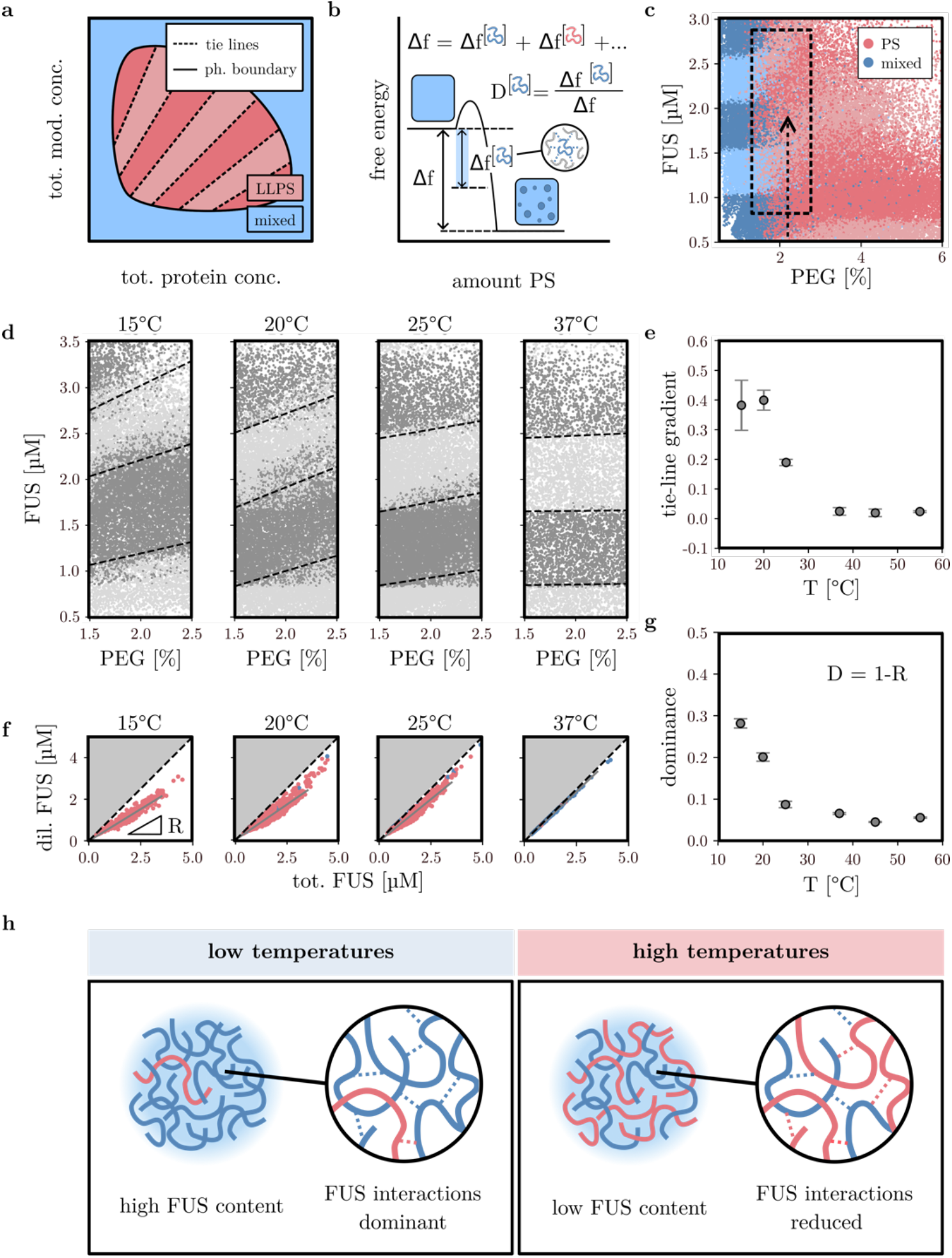
Component stoichiometries and solute free energy contributions to phase separation of the FUS/PEG system at different temperatures. **a** Schematic depicting tie-lines as a line vector connecting protein dense and dilute phase concentrations in protein liquid-liquid phase separation. **b** Schematic illustrating protein dominance D as the free energy contribution of one solute in regard to the total free energy of phase separation. **c** FUS phase diagram as a function of PEG at 25°C. The homogeneous regime is shown in blue, the phase separated regime in red. N = 66311 data points. Protein dilute phase analysis of droplets yields corresponding dilute phase concentrations for each phase separated droplet. The light and dark bands in the phase diagram represent bands of dilute phase concentrations with a < dil. phase conc. < b. The box shows the region next to the phase boundary which is selected for tie-line gradient analysis. The arrow indicates the position of the phase diagram from which 1D line scan data is extracted. **d** Regions between 1.5 to 2.5% PEG were selected from the phase diagram at each temperature for tie-line analysis. Tie-lines are fitted as boundaries between dilute phase bands. For visualization, dilute phase bands with a width of 1µM are shown in different shades of grey. Tie-line fitting was performed using linear SVM. N = 15866, 16304, 19274, 7566. **e** Tie-line gradients as a function of temperature. Average gradients [µM FUS/%PEG] are shown for tie-lines at 1, 2 and 3µM dilute FUS concentration. **f** 1D line scan data at a fixed PEG concentration extracted from the PhaseScan measurement can be used to analyse the FUS dilute phase concentration as a function of the total FUS concentration. Phase separated droplets are shown in blue, homogeneous droplets in red. The dotted diagonal line indicates the case FUS dil. = FUS tot. conc. Dilute phase plots at 3% PEG are shown for 15, 25 and 37°C, respectively. N = 648, 912, 271. **g** Protein dominance as a function of temperature. Average FUS dominance was calculated from 1D line scans between 2 and 2.5% PEG. **h** Schematic illustrating the changes of condensate stoichiometry and FUS interactions with changing temperature.

Using our microfluidic platform, we measured dilute phase concentrations of FUS in droplets over the two-dimensional chemical space to determine the approximate tie-line gradient change with temperature in the low PEG regime (Fig. 2c box with dashed lines, Fig. 2d) using a previously described method^44^. In short, tie-line gradients at temperatures between 15 and 55°C were determined by plotting a phase diagram with dilute phase bands with a FUS concentration between a < [FUS dil.] < b followed by linear fitting of the dilute phase boundaries. These boundaries correspond to high-dimensional tie-line vectors reduced to the measurement plane^47^. We find that with increasing temperature, the tie-line gradient decreased from 0.4 to 0 (Fig. 2e), indicating that the stoichiometry changed towards a lower FUS to PEG ratio inside the condensates.

We measured the dominance of FUS at different temperatures to investigate if change in free energy contribution coming from protein interactions can explain the observed changes in phase behaviour. From phase diagrams, we extracted 1D data slices at fixed PEG concentrations between 2 and 2.5% PEG (Fig.2c, black dashed arrow), and analysed FUS dilute phase concentration as a function of the total FUS concentration at different temperatures (Fig.2f). The data shows that upon temperature increase, the FUS content in the dilute phase increases. At lower temperatures, the gradient of the dilute phase as a function of the total FUS concentration is < 0. With increasing temperature, the gradient increases, a larger fraction of the total protein concentration is found in the dilute phase. At 37°C the dilute phase concentration of FUS becomes equal the total FUS concentration (the gradient R equals 1), i.e. - the majority of the condensates have dissolved.

By fitting a line to the data, we determined the response gradient R of the dilute phase plot which enabled us to estimate the protein dominance parameter D as D = 1-R, a quantitative measure of the contribution of FUS interactions to the overall free energy of FUS/PEG phase separation. Between 15°C and 55°C, the protein dominance decreases significantly from 0.28 to 0.06. The data suggest that with increasing temperature the contributions of protein interactions to the free energy of phase separation decrease. Above 37°C, the protein dominance reaches a constant minimal level between 0 and 0.05. These results can be rationalised by a higher entropic free energy gain of mixing at higher temperatures, leading to an effective weakening of protein interactions and ultimately condensate dissolution.

### Temperature induced changes in protein interactions shed light on RNA recruitment to G3BP1 condensates

Stress granules are involved in the cellular heat stress response. Upon heat stress, the translation initiation factor eIF2*α* is phosphorylated and the cell releases untranslated mRNA from polysomes. This triggers the formation of SGs, membrane-less organelles composed of RNA-binding proteins, small ribosomal subunits and mRNA which have the function to sequester and protect the RNA from damage^63–65^. One of the scaffold proteins which are required for the formation of SGs is G3BP1. It was demonstrated that binding of unstructured RNA to G3BP1 is essential for a conformational change from its autoinhibited to functional state which allows the formation of a condensate network^66,67^. However, how RNA is recruited to SGs and how SG composition is regulated at elevated temperatures under heat stress remains elusive. To gain new mechanistic insights into stress granule assembly, we investigate the heterogeneous G3BP1-PolyA system *in vitro* at various temperatures.

First, we acquired phase diagrams of G3BP1 as a function of PolyA concentration at 15-45°C temperature range (Fig. 3a-b). Surprisingly, the change in the temperature of the environment had little effect on the position of the phase boundary of the G3BP1/PolyA system. We hypothesise that entropically favoured attractions outcompete thermal dissolution in this system, driving phase separation at higher temperatures.

**Fig. 3:**
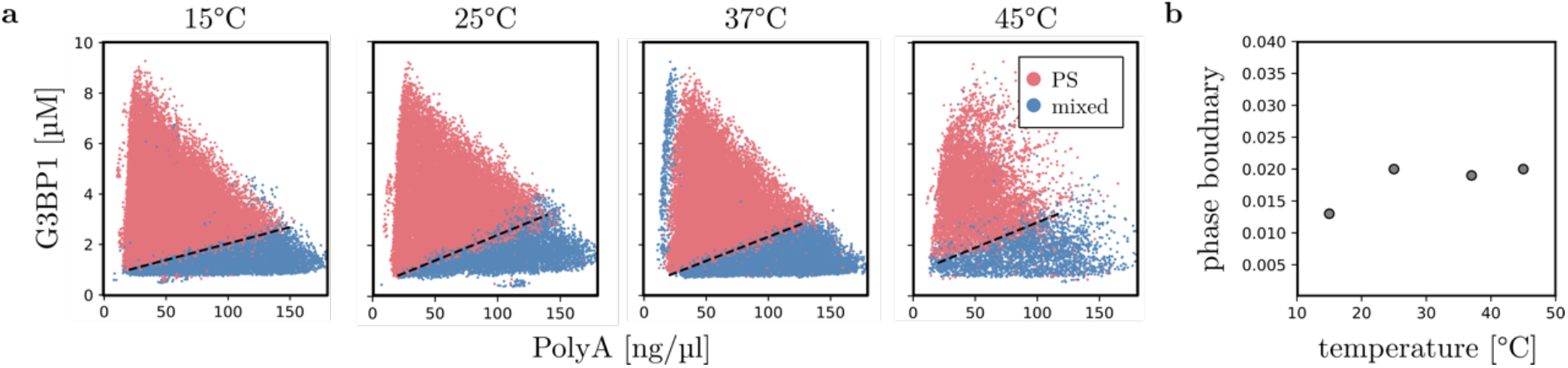
Phase separation of the G3BP1/PolyA system at varying temperatures. **a** Phase diagrams of G3BP1 as a function of PolyA concentration. Points in the phase diagram correspond to individual droplets. The phase separated regime is represented in red, the re-entrant homogeneous phase is shown in blue. N = 34068, 27209, 28788, 11606 data points for 15 to 45°C, respectively. The apparent phase boundaries are shown as a black dotted line. **b** Phase boundary gradients as a function of temperature.

Interestingly, despite the relatively small shift in the phase boundary, we observe a significant change in the gradient of tie-lines (Fig. 4a-b). Between 15°C and 25°C, the tie-line gradient does not change much, above 25°C, however, decreases significantly (Fig. 4c). The data suggest that protein to PolyA ratio remains at a similar level until 25°C and only decreases at elevated temperatures. Additionally, we find that the response gradients R of dilute phase to total protein concentration increase (Fig. 4d).

**Fig. 4:**
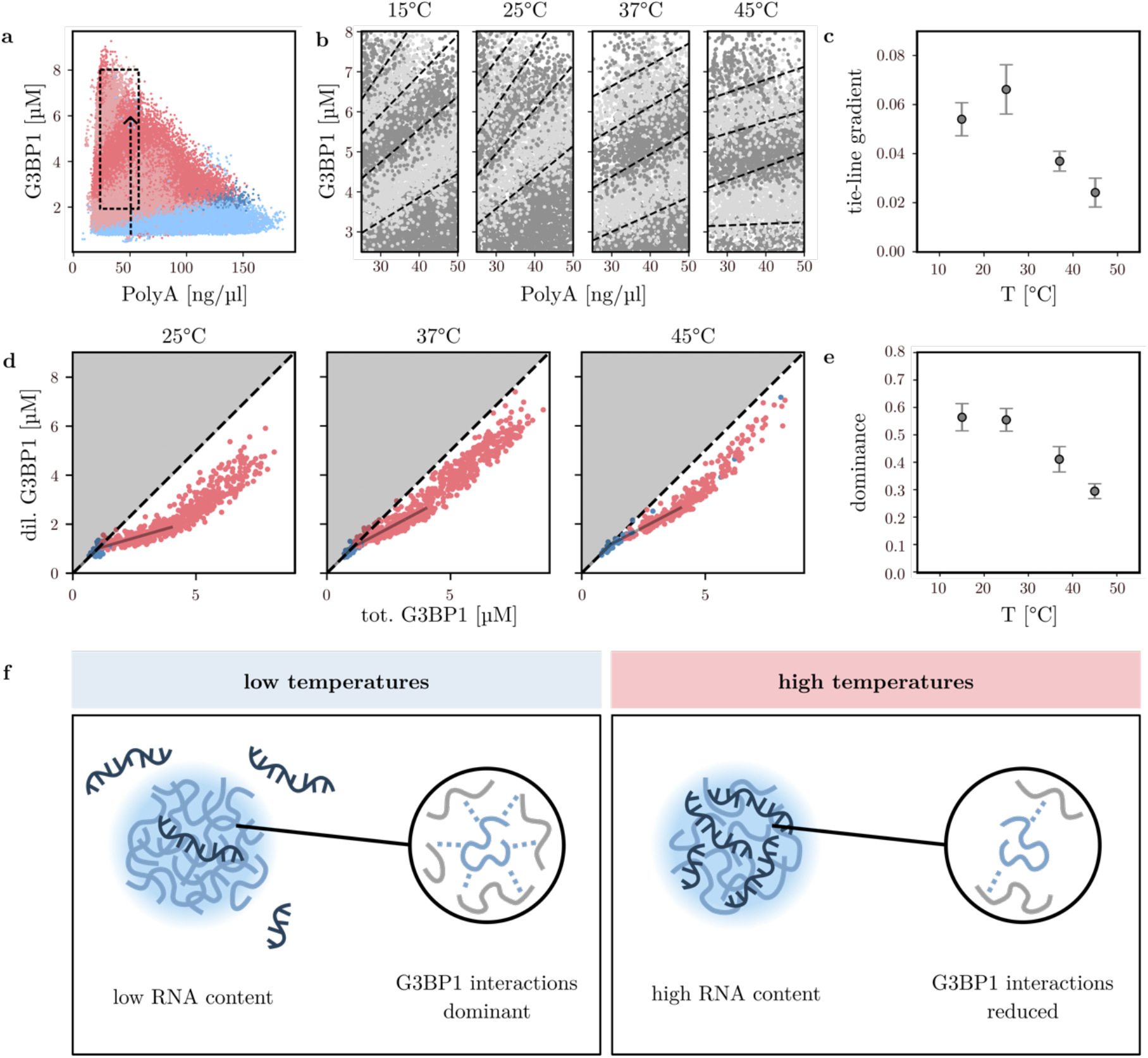
Component stoichiometries and solute free energy contributions to phase separation of the G3BP1/PolyA system at varying temperatures. **a** Phase diagram of G3BP1 as a function of PolyA at 15°C. Phase separated droplets are shown in red, mixed droplets in blue. Dilute phase bands with a width of 2µM G3BP1 are shown as dark and light bands in the phase diagram. The box indicates the region used for tie-line gradient analysis. N = 34068. **b** Dilute phase bands for the G3BP1-PolyA system with a width of 1.2µM are shown in light and dark shades. Tie-lines are fitted as boundaries of dilute phase boundaries using linear SVM. **c** Tie-line gradients [µM G3BP1/% PEG] shown as a function of temperature. N = 9480, 7681, 7524, 5860. **d** Dilute phase plots as a function of total protein concentration at 15, 25, 37 and 45°C. 1D-line scans were extracted at 40ng/µl PolyA. N = 658, 492, 691, 449. **e** Protein dominance as a function of temperature. **f** At low temperatures, G3BP1 interactions dominate the free energy gain of phase separation, the RNA content in condensates is low. At higher temperatures, G3BP1 interactions are reduced and compensated for by other solute interactions, RNA is recruited into the dense phase.

Consequently, the dominance of G3BP1 and therefore the contribution of G3BP1 interactions to the overall free energy of phase separation decreases (Fig. 4e). We hypothesised that entropically favoured interactions involving RNA^68^ could be driving phase separation (Fig. 4f). To validate this, we find that 1,6-hexanediol and urea can dissolve G3BP1-RNA condensates (Supplementary Fig. S14), suggesting that hydrophobic interactions are involved in phase separation of the G3BP1-PolyA system. An increase in hydrophobic interactions with temperature by a favourable increase in desolvation entropy could compensate for loss of G3BP1 dominance, explaining why the phase boundary does not shift.

We interpret our findings in the context of RNA recruitment to SGs as a cytoprotective mechanism during heat shock (Fig. 4f). We propose that as temperature increases to temperatures higher than the physiological level, RNA interactions become favoured over G3BP1 interactions, leading to the recruitment of untranslated RNAs into SGs where they are protected from damage. When the cell returns to normal conditions, RNA is released back into the cytosol, in accordance with our findings: a decrease in tie-line gradients at lower temperatures.

## Conclusion

We showed that on a molecular level, the mechanism of temperature modulation of protein phase separation can be explained through changes in protein interactions. Using our microfluidic approach, we acquire protein phase diagrams and quantify changes in the phase boundary as well as collective, emergent interactions as a function of temperature. By measuring protein dilute phases, we can determine tie-line shifts indicating stoichiometric changes in the dense phase as well as protein dominance, quantifying the relative contribution of protein interactions to the free energy of phase separation. For a typical UCST system such as FUS/PEG, we find a stoichiometric change towards less protein in the dense phase, alongside a reduction of protein interactions with increasing temperature, leading to thermal dissolution of protein condensates.

Further, we investigated how temperature may affect SG stability, composition and function. We used a minimal *in vitro* model system consisting of the SG scaffold protein G3BP1 and unstructured PolyA-RNA. Surprisingly, we find that the phase boundary between the mixed and phase separated regime for the heterogeneous G3BP1/PolyA system is only marginally affected by increasing temperatures. This suggests that in the context of SG formation under heat shock, thermal stability of condensates could be essential for their proper function, ensuring protection of untranslated RNAs inside the stress granule until the cell returns to normal conditions. Interestingly, despite the only small changes in the phase boundary, the composition of the G3BP1/PolyA condensates changes significantly. We find that at temperatures above room temperature, the tie-line gradients of the system as well as the protein dominance decrease, indicating a change in condensate stoichiometry towards a higher RNA content as well as a reduced contribution of G3BP1 interactions to phase separation, compensated for by higher RNA hydrophobic interactions. In the context of heat SG formation, we propose that temperature could regulate the composition of stress granules by finetuning solute interactions, facilitating RNA recruitment to the stress granule during heat stress while its stability is retained.

Overall, we show that our approach can be used to quantify temperature-modulation of phase separation for a variety of systems. By measuring changes in the phase boundary with temperature, alongside with condensate stoichiometries and protein interactions, we can uncover mechanisms of temperature modulation on the molecular level and link them to their observable macromolecular effects on phase separation. We anticipate that our method opens the possibilities of investigating the mechanisms of temperature-modulation of disease-linked phase separating protein systems *in vitro* as well as understanding their effects with a single method, advancing therapeutic developments.

## Acknowledgements

The research leading to these results has received funding from the European Research Council under the European Union’s Seventh Horizon 2020 research and innovation program through the ERC grant DiProPhys (agreement ID 101001615, C.M.F.), Transition Bio (H.A., D.Q., R.S., S.Q.), the Alzheimer Association Zenith Award (P.S.G.-H.).

## Author contributions

C.M.F. H.A., T.S., and T.P.J.K. designed and conceptualised the study. C.M.F. and T.S. performed experiments. S.Q., P.S.G.-H., G.K., and T.P.J.K. provided materials and methods. C.M.F., D.Q., H.A., T.S., R.S. and T.P.J.K. analysed and interpreted the data. C.M.F., H.A. and T.S. wrote the original draft of the paper. All authors discussed the results, commented on the manuscript, and contributed to the final manuscript.

## Correspondence

Correspondence and requests for materials should be addressed to Tuomas P. J. Knowles.

## Data availability

The raw data and analysis code underlying this study will be made available upon request.

