## Supplementary Information for "Temperature-induced changes in protein interactions control RNA recruitment to G3BP1 condensates"

### Materials and Methods

#### Protein expression

Recombinant FUS-GFP was produced according to Qamar et al.[1]
Briefly, recombinant FUS-GFP was expressed in Sf9 insect cells with a baculovirus system. A maltose-binding protein (MBP) tag and a histidine tag (His) were used for protein purification. The infected Sf9 cells with the His-FUS-GFP-MBP construct were grown for 72h after infection. For harvesting, they were centrifuged at 4000rpm for 30 min and resuspended in 50mM Tris HCl pH 7.4 (ThermoFisher Scientific), 500mM KCl (ThermoFisher Scientific), 0.1% CHAPS (ThermoFisher Scientific), 5% w/v glycerol (Merck) and 1mM DTT (Merck). Proteins were purified by Ni-NTA, Amylose affinity column and size exclusion chromatography in the same buffer.

Recombinant G3BP1-emerald was produced according to Qian et al.[2]
Briefly, recombinant G3BP1-emerald was expressed in Sf9 insect cells with a baculovirus system. A maltose-binding protein (MBP) tag and a histidine tag (His) were used for protein purification. The infected Sf9 cells with the His-Emerald-G3BP1-MBP construct were grown for 72h after infection. For harvesting, they were centrifuged at 4000rpm for 30 min, the lysate was further clarified by centrifugation at 10 000G and resuspended in 50mM Tris HCl pH 7.4, 1M KCl, 1mM EDTA 0.1% CHAPS with added protease inhibitor cocktail. Proteins were purified by Ni-NTA, Amylose affinity column and size exclusion chromatography in 50mM Tris pH 7.4, 300mM KCl, 1mM DTT.

#### Device design and fabrication

Microfluidic devices for PhaseScan were fabricated using a photolithographic method. The devices were designed using AutoCAD software by AutoDesk, USA, and printed onto micro-lithography acetate masks by Micro Lithography Services, UK. These masks were then utilised to fabricate a SU-8 mould from which microfluidic devices could be templated. For that, SU-8 would be poured onto a silicon wafer an equally distributed by spinning the wafer at 500-3000 rpm for 45 s to the desired height of 50µm. Then, the wafer would be pre-baked on a heat plate for 5-10 min at 95°C. After pre-baking, the acetate mask was placed on top of the wafer and the wafer was illuminated under UV light at 365nm for 60s to crosslink the SU-8 polymer at the exposed sites not covered by the mask. After UV exposure, the wafer was post-baked for 5 min at 95 °C to harden the polymer cross-links. Removal of non-crosslinked SU-8 was achieved by washing the wafer with propylene glycol monomethyl ether (PGMEA, Sigma-Aldrich) and isopropyl alcohol (IPA, Sigma-Aldrich). The height of the mould was finally determined using a profilometer by Dektak, Bruker, USA. To produce the desired microfluidic devices, the wafer was placed in a petri dish and covered with a freshly prepared mixture of liquid poly-dimethylsiloxane (PDMS) and Sylgard184 by Dow

Corning. The PDMS was degassed for 30 min in a desiccator, then polymerised by baking at 65 °C for 90 min. After baking, the PDMS device was cut out of the petri dish and holes were punched into the PDMS at the inlet and outlet positions. After cleaning by sonicating the device in an isopropyl alcohol bath for 15 mins and drying it for 1h at 65°C, the device and a microscope slide were treated in an oxygen plasma oven by Electronic Diener at 40 % power for 45s, Then the device was placed on top of the slide and slightly pressed together for bonding. Finally, the channels of the microfluidic devices were treated with a mixture of hydrofluoroether (HFE-7500) (Fluorochem) and 1.5 % trichloro(1H,1H,2H,2H- perfluorooctyl)silane (Sigma-Aldrich).

**Protein sample preparation, droplet making and temperature control**

Experiments were performed under physiological conditions, at 150mM KCl and in 50mM Tris HCl pH 7.4. For experiments with G3BP1, 2 wt/v% PEG 10K (ThermoFisher Scientific) was added to all buffers. A protein solution of 5µM FUS-GFP or 10µM G3BP1-emerald in physiological buffer was prepared by diluting the protein from a protein stock in 1M KCl, 50mM Tris pH7.4. PEG solutions for experiments with FUS were prepared by dissolution of 10 wt/v% PEG 10K in 50mM Tris HCl pH7.4, 150mM KCl and barcoded with 3µM free AF647 carboxylic acid (ThermoFisher Scientific) to be able to determine the PEG concentrations in the droplets. For G3BP1 experiments, solutions of 200ng/µl PolyA RNA (Merck) in physiological buffer were prepared and barcoded with 3µM AF647 for concentration determination. For the microfluidic experiment, three aqueous solutions containing protein, buffer and modulator (PEG or Poly(A), respectively) as well as an oil solution for droplet generation (HFE-7500 mechanical oil containing 1.2% Bio-RAN) were used. The solutions were loaded into four separate inlets on the microfluidic chip using pressure control pumps (LineUp Flow EZ, Fluigent). The three aqueous solutions were mixed in one channel before reaching the droplet junction, where droplets were generated by oil flow at 100µl/h. By variation of the flow rates of the three aqueous solutions between 5 and 50µl/h at a constant total flow rate of 60µl/h, droplets with the same size but different concentrations of protein and modulator were generated. The droplets were incubated for 4 mins on the microfluidic chip while flowing through the incubation channel until reaching the wider imaging chamber, where they slowed down and were imaged. As the microfluidic chip was positioned on a temperature control stage (Instec TS102Si) for the duration of the experiment, the droplets would undergo incubation at the desired temperature for 4 mins before being imaged.

**Droplet imaging**

Droplets were imaged in continuous flow through the microfluidic chip every 4 seconds using an openFrame epifluorescent microscope (Cairn Research) equipped with a 10x air objective (Nikon CFI Plan Fluor) and a dichroic filter set (Cairn Research) to image three wavelengths (488nm, 546nm, 647nm) at the same time. Crosstalk calibration images were acquired by flowing droplets containing only one dye through the chip, the fluorescence observed in the other two channels was used to calculate the crosstalk.

**Data analysis: droplet detection and phase diagram generation**

Microscopic images of phase separated and homogeneous droplets were analysed using a custom-written Python (Python version 3.9.7) script. The droplets were identified by circle detection and filtered from erroneous droplets by shape and radius. For each wavelength, the fluorescence intensity after subtraction of the illumination background was mapped to an intensity/concentration fit defined as a straight line from the $1^{st}$ to the $99^{th}$ percentile of fluorescence intensity against 0 to stock concentration. The intensity to concentration calibration was performed separately for each temperature to ensure that reduced dye efficiency at higher temperatures is accounted for (Supplementary Fig. 8). Classification as phase separated or homogeneous was performed by a convolutional neural network trained on human annotated data (Supplementary Fig. 12). Phase diagrams were plotted. One datapoint in the phase diagram corresponds to one droplet.

**Data analysis: Tie-line analysis**

Dilute phase concentrations of the proteins were determined using the darkest 5-25% of the pixels in each droplet which were converted to concentrations by the same method as above. Dilute phase bands with concentrations between $a < [\text{protein dil.}] < b$ were fitted using a linear SVM fit to yield the tie line gradient.

**Data analysis: Protein dominance**

1D line scan data at a fixed modulator (PEG for FUS, PolyA for G3BP1) concentration was extracted for dilute phase plots at different temperatures. Plots of protein dilute phase as a function of the total protein concentration were used to calculate protein dominance. The line scan data was fitted using a linear fit to determine the gradient R of the dilute phase as a function of the total phase. The dominance, i.e. the free energy contribution of the protein to the total free energy of phase separation, was calculated using the relationship $D = 1\text{-}R$.

**Fluorescence imaging**

Imaging of G3BP1/PolyA condensates was performed using an openFrame epifluorescent microscope (Cairn Research) equipped with a 20x air objective (Nikon CFI TU Plan Epi) and a dichroic filter set (Cairn Research). Images were acquired using µManager software by placing a 5µl aliquot of the sample on a microscope slide on the microscope stage.

### Supplementary Figures

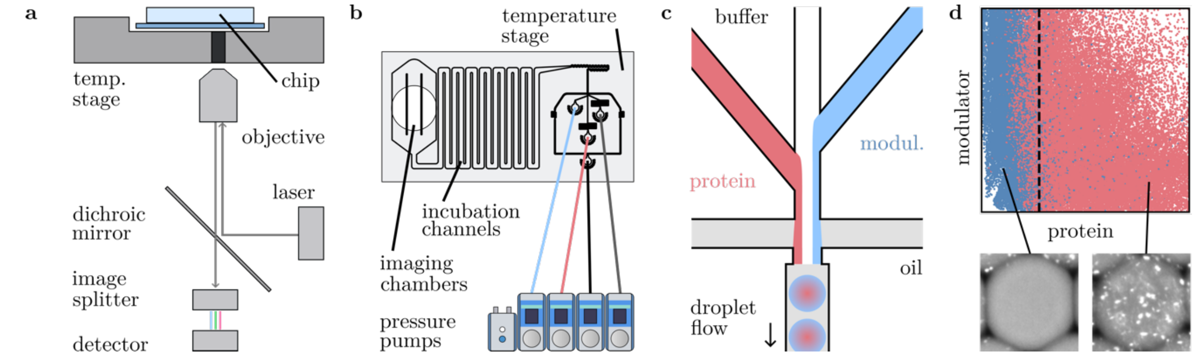


Supplementary Figure S1. A microfluidic chip mounted on a temperature control stage is used to perform the experiments. A standard epifluorescence imaging setup **a** is used for data acquisition, the microfluidics are controlled using pressure pumps **b** and the chip design itself encompasses the standard droplet maker **c**, incubation channels, and large imaging chambers. **d** Condensates are detected in each droplet, and total concentrations of the protein and modulator are determined from total fluorescent intensities. A phase diagram is then plotted using the droplet data.

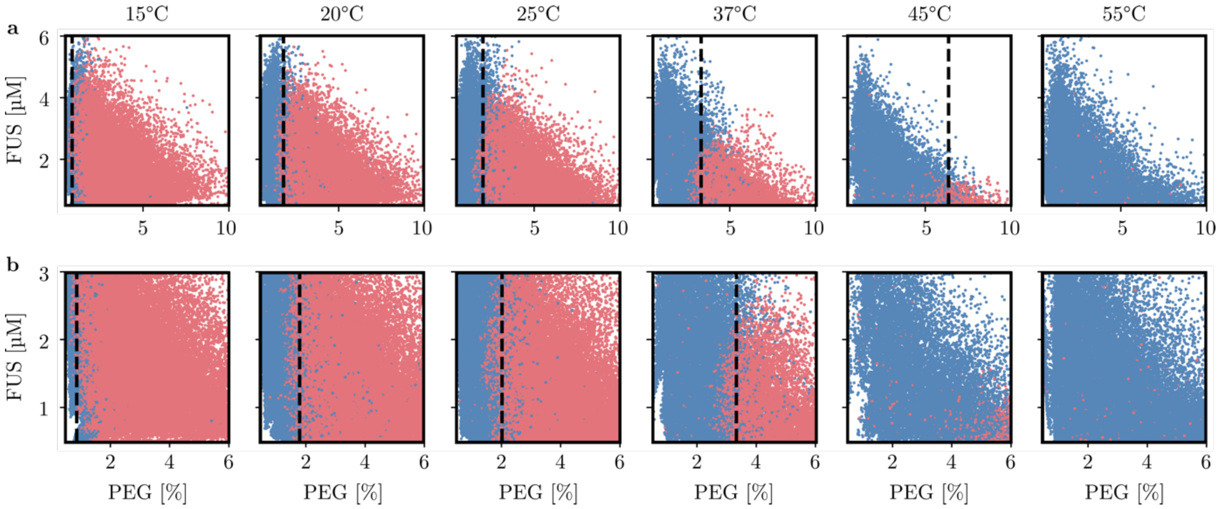


Supplementary Figure S2. **a** Full FUS-PEG phase diagrams at temperatures between 15 and 55°C. In the experiment, FUS concentrations were varied between 0 and 6μM and PEG concentrations were varied between 0 and 10 % PEG 10K. N = 61331, 66311, 74748, 30186, 18396, 47888. **b** Close up of region of interest of FUS-PEG phase diagrams. Phase boundaries were fitted using a linear support vector machine fit.

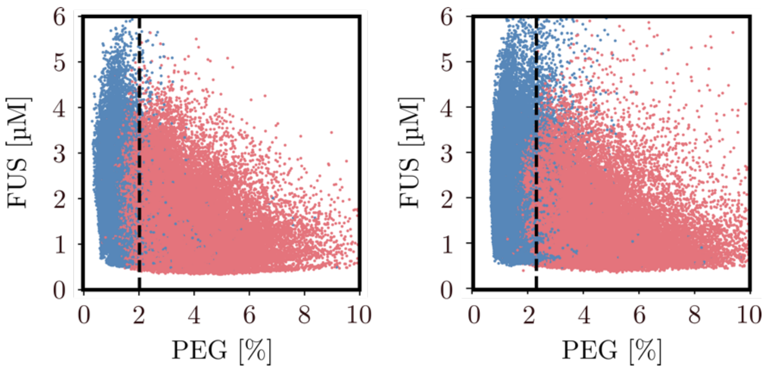


Supplementary Figure S3. Robustness of phase diagram measurements. Phase diagrams of FUS-PEG at 25°C acquired on two different days, using different samples and microfluidic chips. The phase boundary for FUS-PEG at 25°C is at 2.16 ± 0.14 % PEG 10K.

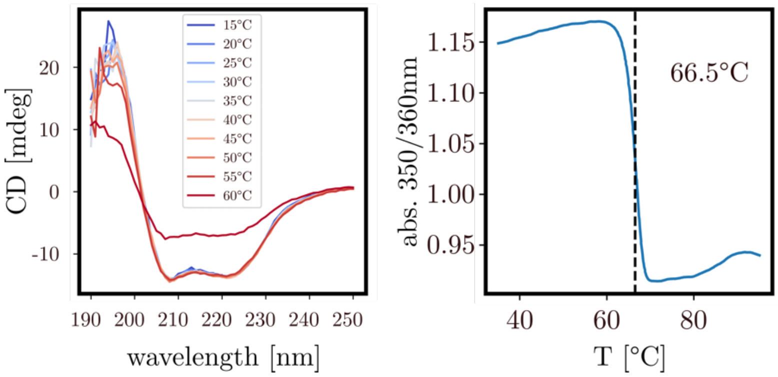


Supplementary Figure S4. Thermal stability of FUS-GFP. Left: CD spectra recorded at temperatures between 15 and 60°C. FUS protein structure is intact at temperatures up to 55°C. Right: UV-vis absorption of FUS-GFP as a function of temperature measured between 35 and 95°C with calculated melting point (dashed line) at 66.5°C.

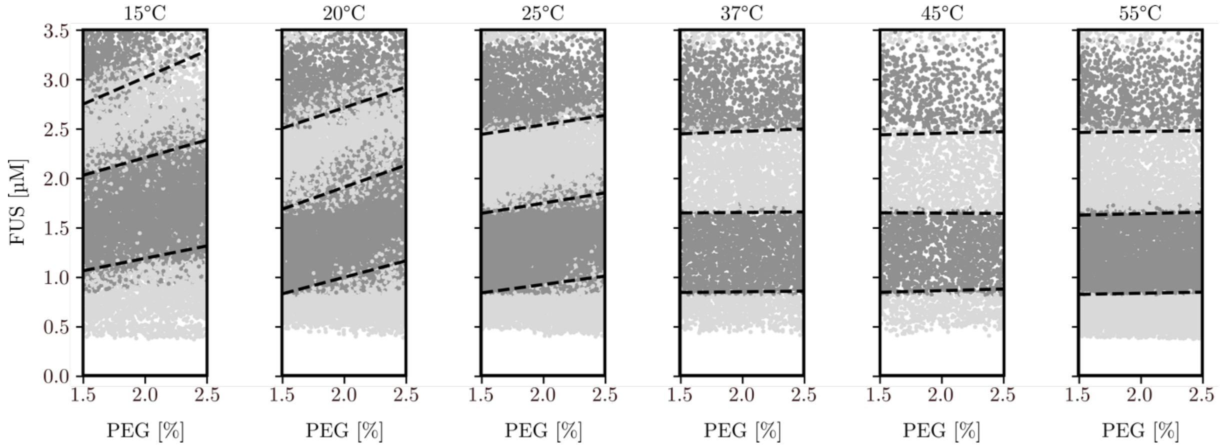


Supplementary Figure S5. Full tie-line plots of FUS-PEG at temperatures between 15 and 55°C. Dilute phase bands with a width of 1μM are plotted in shades of grey. Dilute phase boundaries were fitted using a linear SVM fit. N= 15866, 16304, 19274, 7566, 4958, 13819.

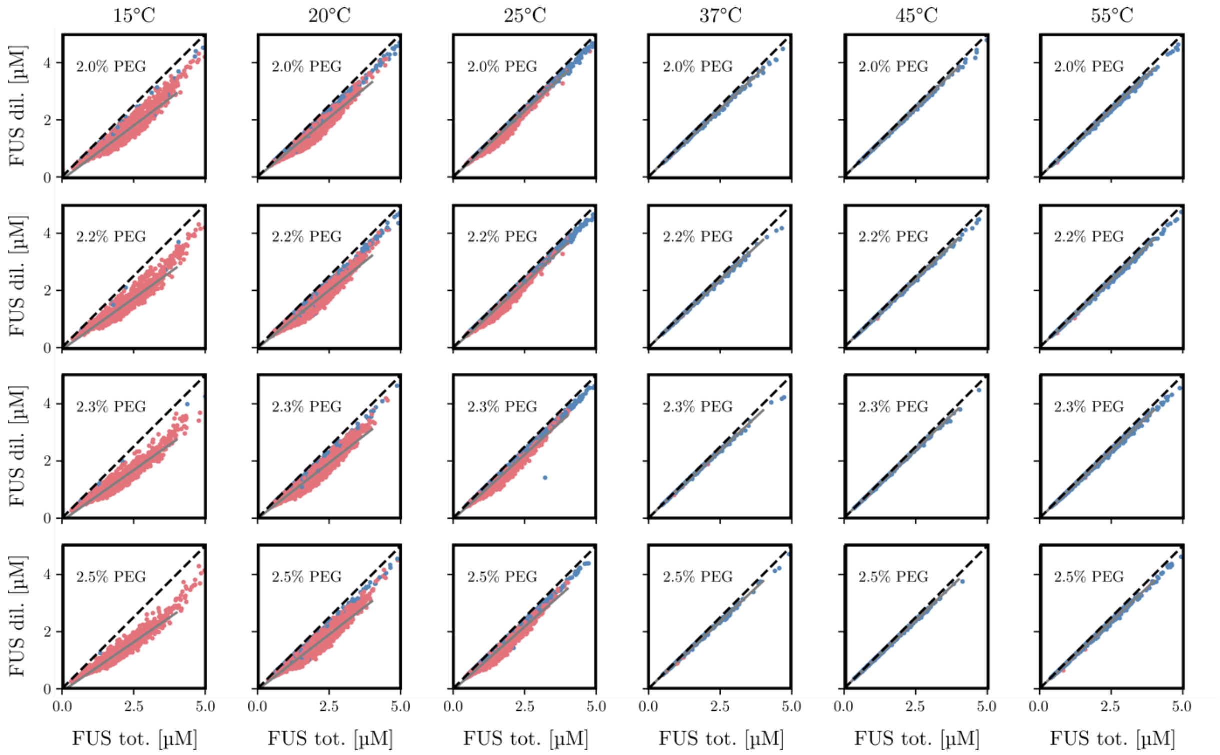

Supplementary Figure S6. Dilute FUS concentration as a function of total FUS concentration at temperatures from 15 to 55°C with 2-2.5% PEG. The response gradient R (grey line) is determined by a linear fit. At low temperatures, the gradient is <1, at high temperatures above 37°C, the gradient is ~1 and the dilute phase concentration equals the total protein concentration, the condensates are dissolved.

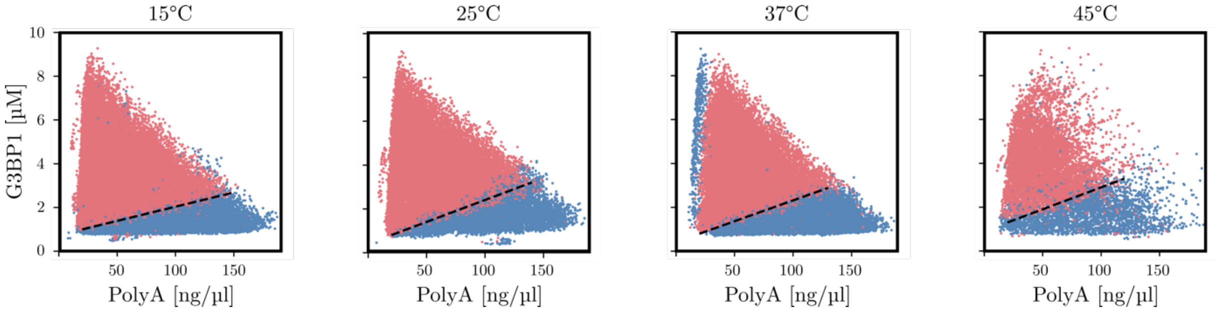


Supplementary Figure S7. Full G3BP1-PolyA phase diagrams at temperatures between 15 and 45°C. In the experiment, G3BP1 concentrations were varied between 0 and 9µM and PolyA concentrations were varied between 0 and 180 ng/µl. N = 34068, 27209, 28788, 16208.

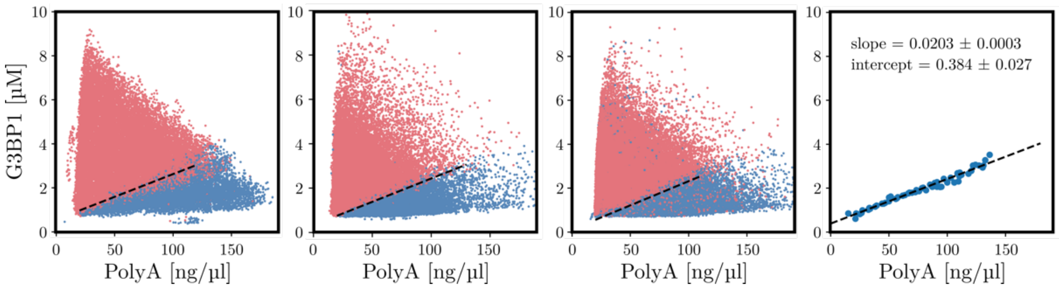


Supplementary Figure S8. Robustness of G3BP1-PolyA phase diagram measurements. Left: Phase diagrams of G3BP1-PolyA at 25°C acquired on three different days, using different samples and microfluidic chips. Right: Superimposed phase boundaries of the three experiments determined by and linear fit. Slope: 0.0203 ± 0.0003, intercept: 0.384 ± 0.027.

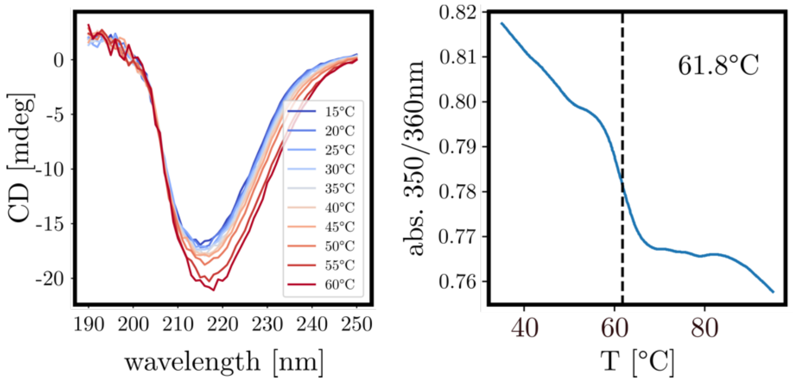


Supplementary Figure S9. Thermal stability of G3BP1-emerald. Left: CD spectra recorded at temperatures between 15 and 60°C. G3BP1 protein structure is intact at temperatures up to 60°C. Right: UV-vis absorption of FUS-GFP as a function of temperature measured between 35 and 95°C with calculated melting point (dashed line) at 61.8°C.

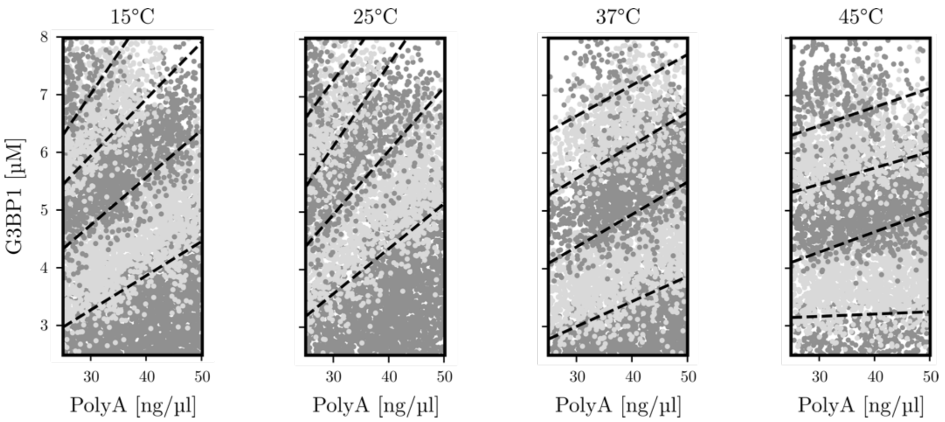


Supplementary Figure S10. Full tie-line plots of G3BP1-PolyA at temperatures between 15 and 45°C. Dilute phase bands with a width of 1.2µM are plotted in shades of grey. Dilute phase boundaries were fitted using a linear SVM fit. N= 9480, 7681, 7524, 5860.

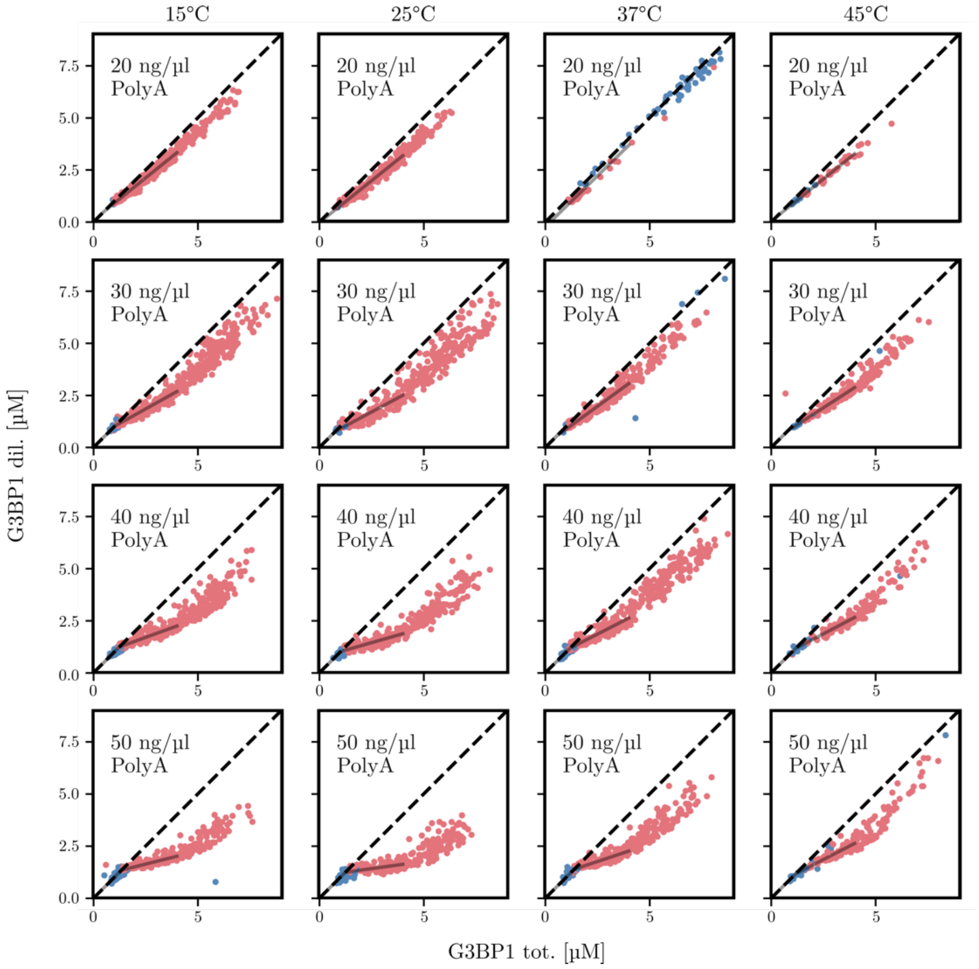


Supplementary Figure S11. Dilute G3BP1 concentration as a function of total G3BP1 concentration at temperatures from 15 to 45°C with 20-50 ng/µl PolyA. The response gradient R (grey line) is determined by a linear fit. At low temperatures, the gradient is significantly <1, at higher temperature, the gradient approximates 1, the dilute phase concentration increases.

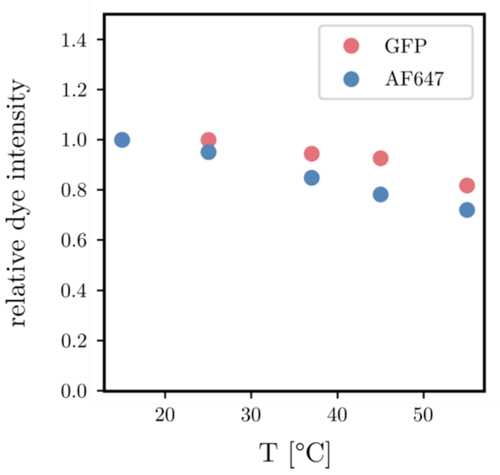


Supplementary Figure S12. Measurement of relative dye intensity as a function of temperature. Dye intensities of droplets containing 3µM GFP or 3µM AF647 were imaged and measured in a temperature range between 15 and 55°C. Fluorescence intensities inside the droplets were compared relative to each other at different temperatures. A decrease in dye signal was observed with increasing temperature. The changes in intensity were accounted for in the droplet analysis of PhaseScan experiments.

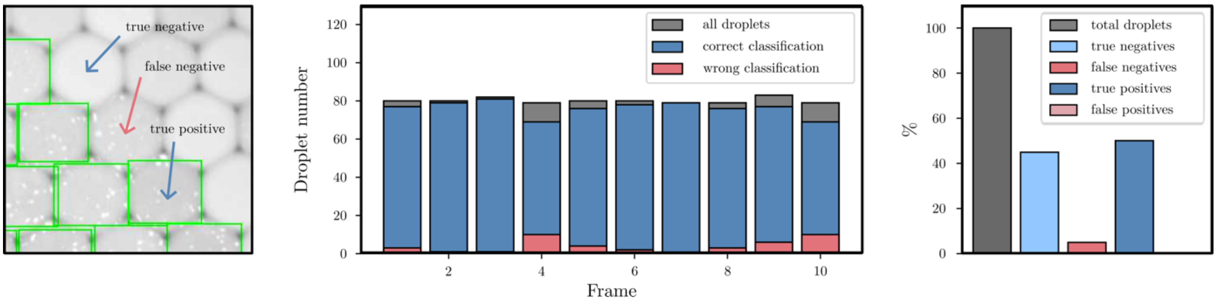


Supplementary Figure S13. a) Example of condensate detection using PhaseScan microscopic images. Droplets classified as phase separated by the CNN are highlighted by a green box. The image shows examples of true negatives (classified mixed droplets), true positives (classified phase separated droplets) and false negatives (phase separated droplets classified as mixed). False positives did not occur. b) Condensate detection over 10 randomly chosen frames of PhaseScan images. Correctly and incorrectly classified droplets are shown as a fraction of the total droplet number in each frame. The number of correctly detected droplets was calculated as $n_{correct} = n_{true\ negative} + n_{true\ positive}$. The number of incorrectly detected droplets was calculated as $n_{wrong} = n_{false\ negatives} + n_{false\ positives}$. c) Relative weights of true and false negative and positive classifications in regard to the total number of droplets detected.

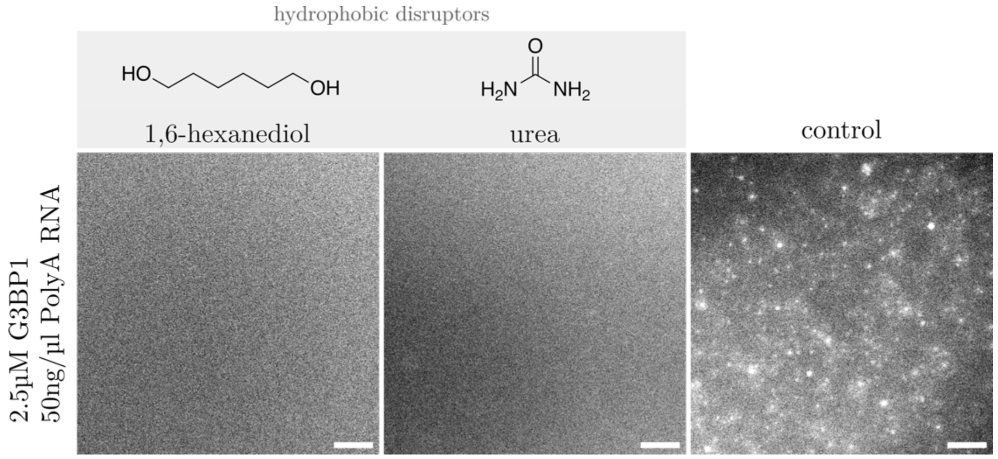


Supplementary Figure S14. Representative microscopic images of G3BP1/PolyA condensates upon addition of the hydrophobic disruptors 1,6-hexanediol and urea. The total concentrations were 2.5µM G3BP1-emerald, 50ng/µl

PolyA RNA and 2% PEG 10K in 150mM KCl, 50mM Tris-HCl pH 7.4. The final additive concentrations were 20% 1,6-hexanediol or 2M urea, respectively. The images were taken at room temperature. The images are representative of the observed reproducible behaviour from at least three test replicates of the respective protein conditions. Scale bar is 100μM.
